# Active zone compaction in presynaptic homeostatic potentiation

**DOI:** 10.1101/802843

**Authors:** Achmed Mrestani, Philip Kollmannsberger, Martin Pauli, Felix Repp, Robert J. Kittel, Jens Eilers, Sören Doose, Markus Sauer, Anna-Leena Sirén, Manfred Heckmann, Mila M. Paul

## Abstract

Brain function relies on neurotransmission which is stabilized by presynaptic homeostatic potentiation (PHP). PHP operates on time scales ranging from minute- to life-long adaptations and likely involves reorganization of presynaptic active zones (AZs). At *Drosophila melanogaster* neuromuscular junctions, earlier work ascribed AZ enlargement by incorporating more Bruchpilot (Brp) scaffold protein a central mechanistic role in PHP.

We used localization microscopy (*direct* stochastic optical reconstruction microscopy, *d*STORM) and hierarchical density-based spatial clustering of applications with noise (HDBSCAN) to study AZ plasticity during PHP. We found that both acute, philanthotoxin (PhTx)-induced and chronic, genetically-induced PHP lead to compaction of individual AZs without altering Brp copy numbers per AZ. This compaction even occurs within Brp subclusters of the AZ scaffold which also move towards AZ centers. Furthermore, lowering imaging resolution revealed how AZ compaction in PHP translates into apparent increases in AZ area and Brp protein content as implied earlier. Our results suggest AZ compaction in PHP as an effective mechanism to raise presynaptic protein density and transmitter release.

**SIGNIFICANCE STATEMENT:** Homeostatic plasticity stabilizes chemical synaptic transmission in multiple organisms ranging from insects to humans. Changes in active zones (AZs), membrane specializations of the presynapse where synaptic vesicles are discharged, are thought to be crucial in homeostatic adaptations. AZ growth by protein incorporation was proposed as a core mechanism in presynaptic homeostatic potentiation (PHP). Localization microscopy of an abundant AZ scaffold protein uncovered that instead of growing, AZs are compacted in acute and chronic PHP. At lower imaging resolution, however, AZs appear larger and brighter although protein numbers are not increased. In summary, our findings suggest AZ compaction as new and effective mechanism to raise presynaptic protein density and transmitter release in PHP.

## INTRODUCTION

Chemical synapses are complex nanomachines that integrate and process information. Synapses are optimized for fast and reliable performance in combination with miniaturization and plasticity (Atwood and Karunanithi, 2002; Chua et al., 2010; Kittel and Heckmann, 2016; Neher and Brose, 2018). An intriguing form of synaptic plasticity is presynaptic homeostatic potentiation (PHP, Turrigiano, 2008; Davis and Müller, 2015). Homeostatic plasticity has been studied at vertebrate synapses (Turrigiano et al., 1998; Murthy et al., 2001; Thiagarajan et al., 2005; Lindskog et al., 2010; Jeans et al., 2017; Delvendahl et al., 2019) as well as at *Drosophila melanogaster* neuromuscular junctions (NMJs, Davis and Goodman, 1998; DiAntonio et al., 1999; Frank et al., 2006; Müller et al., 2012; Younger et al., 2013; Orr et al., 2017; Gratz et al., 2019). While the structural correlates of PHP remain elusive, several mechanisms are conceivable including incorporation, removal, modification and/or rearrangement of key proteins at active zones (AZs), presynaptic membrane specializations where synaptic vesicles are discharged.

The cytomatrix at the active zone (CAZ) consists of an organized set of proteins (Südhof 2012). Large α-helical coiled-coil proteins of the ELKS/CAST family are crucial for the regulation of synaptic transmission (Held et al., 2016; Dong et al., 2018). In *Drosophila*, the ELKS/CAST homolog Bruchpilot (Brp) is essential for synchronous glutamate release and its amount was shown to correlate with release probability and structural synaptic differentiation (Kittel et al., 2006; Ehmann et al., 2014; Peled et al., 2014; Paul et al., 2015; Akbergenova et al., 2018). Brp is a major AZ scaffold protein responsible for clustering of presynaptic calcium channels close to release sites creating a proper molecular environment for fast and precise neurotransmitter release (Kittel et al., 2006; Held and Kaeser, 2018). Whereas the Brp N-term was mapped in membrane-proximity, its C-term localizes about 155 nm above the postsynaptic receptors (Fouquet et al., 2009; Liu et al., 2011) and is important for tethering synaptic vesicles (Hallermann et al., 2010). Remarkably, Brp is distributed heterogeneously at *Drosophila* NMJs forming about 15 subclusters within an AZ (Ehmann et al., 2014). Subclusters with high protein concentration should promote vesicle tethering as shown for Syntaxin (Sieber et al., 2007; Barg et al., 2010; van den Bogaart et al., 2011). Early work at *Drosophila* NMJs showed no evidence for protein synthesis during PHP (Frank et al., 2006). However, confocal and STED microscopy suggested an increase in Brp amount during acute and chronic PHP (Weyhersmüller et al., 2011; Goel et al., 2017; Böhme et al., 2019).

Here, we reason that localization microscopy in terms of *direct* stochastic optical reconstruction microscopy (*d*STORM, Heilemann et al., 2008; van de Linde et al., 2011; Löschberger et al., 2012) permits precise quantification of Brp in PHP (Ehmann et al., 2014; Paul et al., 2015). For unbiased data analysis we perform hierarchical density-based spatial clustering of applications with noise (HDBSCAN, Campello et al., 2013). We address the following fundamental questions: How robust are localization-based results relying on density-based clustering algorithms? Does the amount of protein within individual AZs increase during PHP and/or does the nanoarchitecture of the AZ scaffold change? Do AZs grow or shrink? Using data simulations and correlative confocal-*d*STORM imaging we analyze the relation between thresholding- and localization-based fluorescence quantification of small structures below the diffraction limit. We discover striking differences of Brp organization during acute, pharmacologically-induced and chronic, genetically-induced homeostasis and reason that AZ compaction and not AZ growth is the structural correlate of release strengthening in PHP.

## RESULTS

### Analysis of super-resolution data

We imaged type Ib boutons on abdominal muscles 6 and 7 of male *Drosophila melanogaster* 3^rd^ instar larvae (Figure 1A) using *d*STORM (Heilemann et al., 2008; van de Linde et al., 2011; Löschberger et al., 2012; Ehmann et al., 2014; Paul et al., 2015) and a highly specific monoclonal antibody Brp^Nc82^, that maps to the C-terminal region of Brp (Kittel et al., 2006; Fouquet et al., 2009). Since Brp is an abundant protein and our epitope in Brp covers the spatial extent of individual AZs, we interpret ‘Brp area’ as ‘AZ area’. Density-based clustering algorithms allow to study synaptic nanotopology (Bar-On et al., 2012; Ehmann et al., 2014; Szoboszlay et al., 2017; Rahbek-Clemmensen et al., 2017; Rebola et al., 2019). We used a Python implementation of hierarchical density-based spatial clustering of applications with noise (HDBSCAN), which robustly extracts clusters in data with varying density (Campello et al., 2013; McInnes et al., 2017). To test the algorithm’s robustness, we varied its free parameters ‘minimum cluster size’ and ‘minimum samples’ and analyzed their influence on the number of detected Brp clusters, the number of Brp localizations and the area per cluster (Figure 1B-E, Supplementary Figure 1). A wide range of parameters delivered robust results and all Brp clusters in the following analyses were extracted with the combination 100 and 25 for minimum cluster size and minimum samples, respectively.

**Figure 1.**
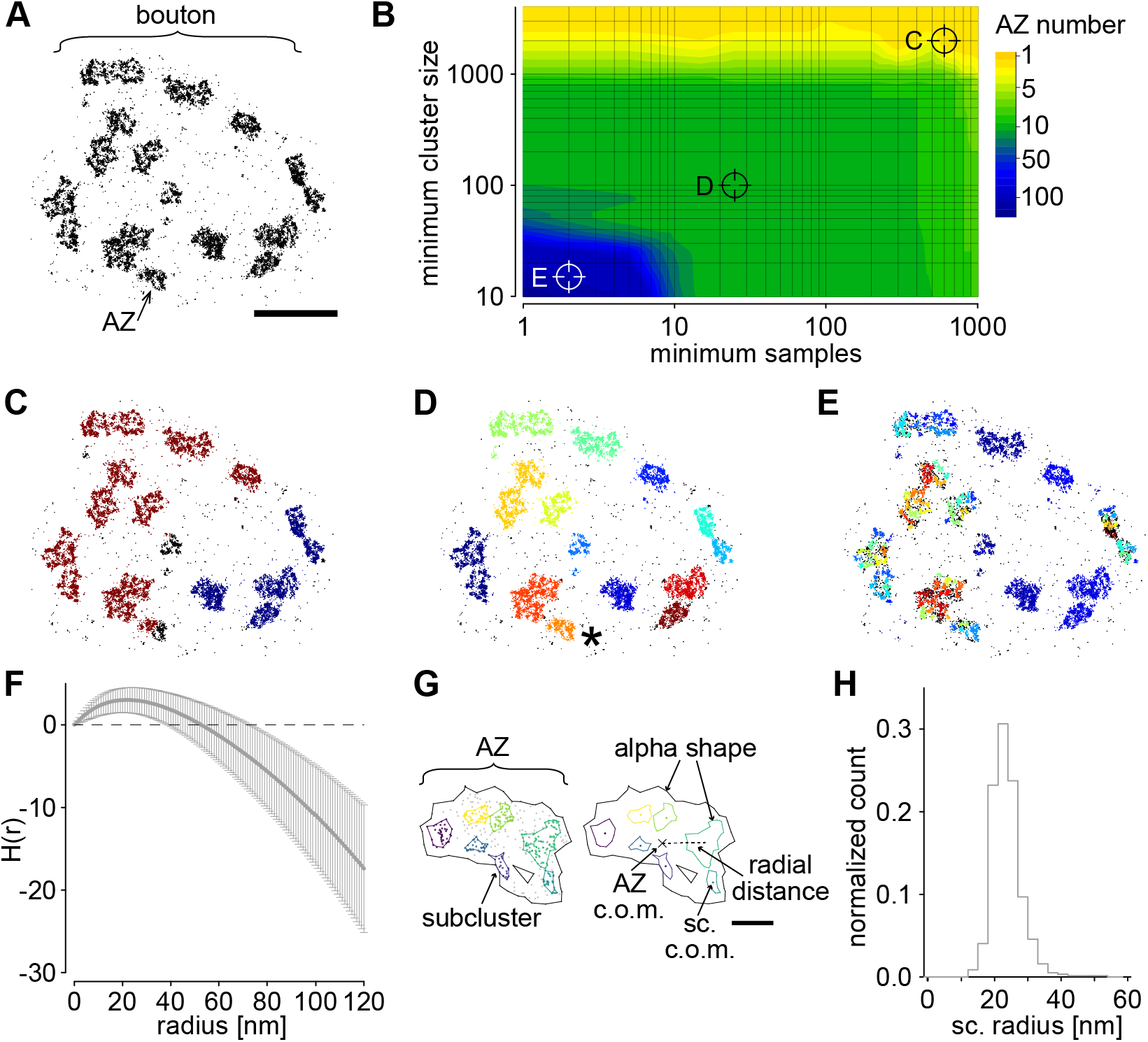
Cluster analysis of presynaptic Brp localizations. (A) Scatter plot of 15470 localizations from *d*STORM of a representative type Ib bouton of a wildtype *Drosophila* NMJ on abdominal muscles 6/7 stained with Brp^Nc82^ antibody labelled with Alexa Fluor647 conjugated F(ab’)_2_ fragments. Arrow indicates an individual cluster of Brp localizations, defined as an AZ throughout the manuscript. (B) Contour plot of the number of detected AZs in the data from (A) depending on the HDBSCAN clustering parameters ‘minimum samples’ and ‘minimum cluster size’. Parameter combinations for clustering in (C-E) are indicated. (C-E) Data from (A) after HDBSCAN analysis. Colors indicate cluster identity, unclustered localizations are displayed in black. Parameter combinations for minimum cluster size and minimum samples were 2000 and 600 (C), 100 and 25 (D) and 15 and 2 (E). Note that multiple AZs are merged into two large clusters in (C) and that individual AZs are split into multiple small clusters in (E). Minimum cluster size 100 and minimum samples 25 are used for AZ detection throughout this manuscript. Asterisk in (D) highlights the AZ cluster shown in (G). (F) Averaged H function (grey, mean ± SD, derivative of Ripley’s K function) from n = 568 AZs obtained from 13 NMJs from 6 wildtype animals. The maximum of the curve indicates a mean subcluster (sc.) radius of 22 nm. Dashed black line indicates the prediction for a random Poisson distribution. (G) HDBSCAN for subcluster detection applied to an individual AZ (asterisk in D). Alpha shapes used for AZ area quantification are indicated by black lines. Left: Colored Brp subclusters surrounded by colored lines indicating alpha shapes. Unclustered Brp localizations are shown as grey dots. Right: The centers of mass (c.o.m.) of this AZ (cross) and of each subcluster (colored dots) are indicated. A dashed line shows the Euclidean distance between the AZ c.o.m and an individual sc. c.o.m., referred to as ‘radial distance’. (H) Histogram of mean Brp subcluster radius per AZ assuming a circular area, median (25^th^-75^th^ percentile): 23 (21-26) nm) for AZs in (F). Scale bars 1 μm in (A) and 100 nm in (G).

To probe the existence of Brp subclusters in our data we computed H functions (derivative of Ripley’s K function) and calculated an averaged curve (Figure 1F). Positive values for H(r) indicate clustering, negative dispersion or edge effects, and maximum positive values roughly correspond to the radius of putative clusters (Kiskowski et al., 2009). The maximum of the curve at 22 nm is in agreement with the dimensions of Brp subclusters reported in earlier work (Ehmann et al., 2014). Subsequently, we performed a second HDBSCAN on individual AZs with adjusted parameters that yielded similar subcluster radii as the H function (Figure 1G, H). In addition to alpha shape generation for area determination, localization-based analysis yields the possibility to define a center of mass (c.o.m.) of an individual subcluster and of the entire AZ and to measure their specific interspaces (Figure 1G). The distance between AZ c.o.m. and subcluster c.o.m. will be referred to as radial distance.

### Compaction of the Bruchpilot scaffold in acute and chronic homeostasis

The wasp venom Philanthotoxin (PhTx) induces PHP within minutes and increases quantal content and release-ready vesicles in our preparation (Frank et al., 2006; Weyhersmüller et al., 2011; Davis and Müller, 2015). We compared AZs of type Ib boutons incubated in PhTx (phtx) or DMSO (control, Figure 2A). Brp localization numbers as a measure for the amount of Brp protein per AZ (Ehmann et al., 2014) were not changed in phtx, however, AZ area was reduced compared to controls and, hence, Brp density increased in phtx (Figure 2B, see Supplementary Table 1 for all data presented here). Thus, PHP induces a rearrangement of Brp without changing the number of Brp molecules within individual AZs of type Ib boutons.

**Figure 2.**
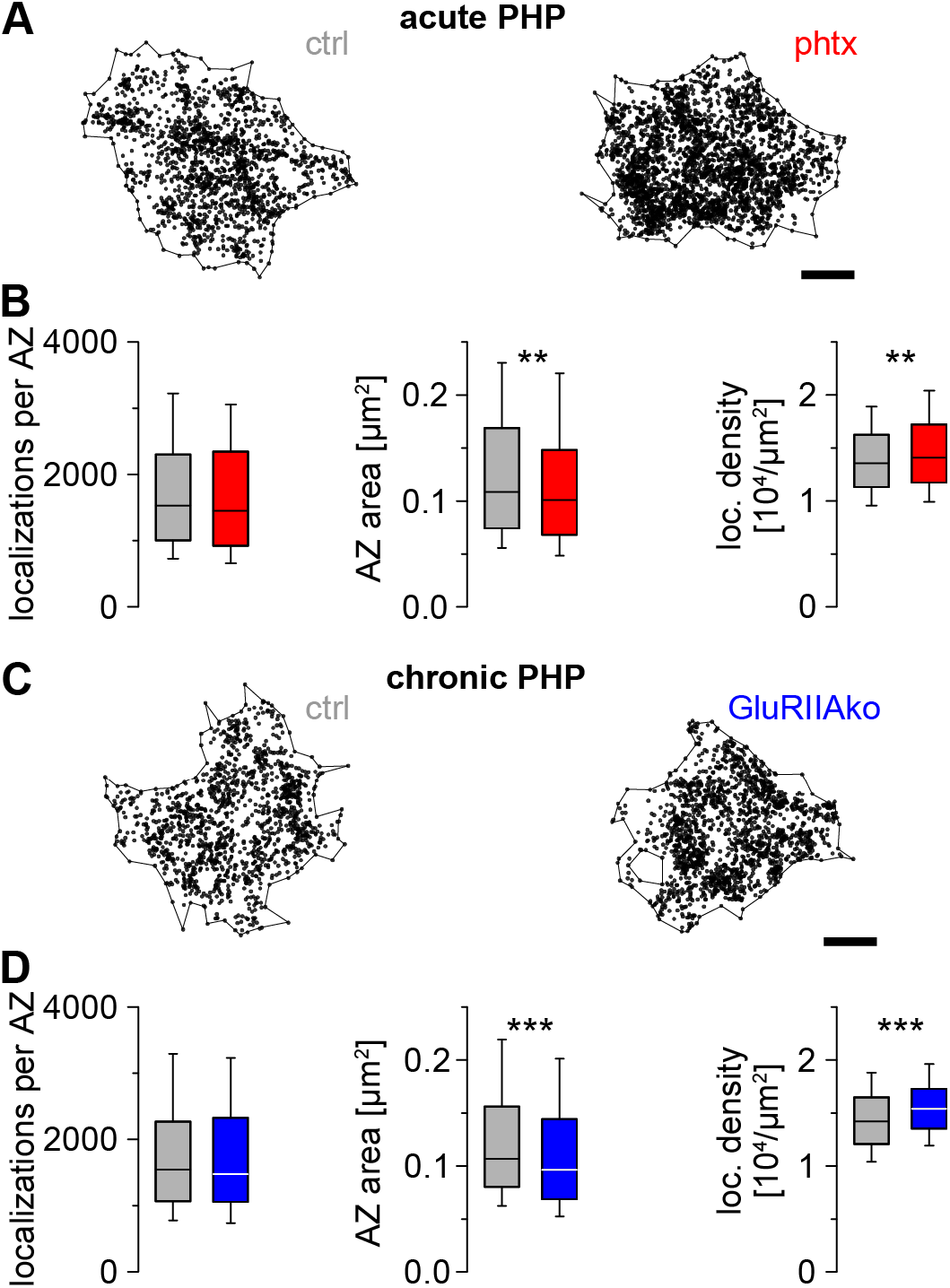
Acute and chronic homeostatic potentiation decrease AZ area and increase Brp density. (A) Scatter plots of type Ib AZs from a control (ctrl, left) and a Philanthotoxin treated animal (phtx, right) for induction of acute presynaptic homeostatic potentiation (PHP). Localizations outside alpha shape are not shown for clarity. (B) Number of Brp localizations per AZ (p = 0.236), AZ area (p = 0.009) and Brp localization density (p = 0.009) for ctrl (grey, n = 568 AZs from 13 NMJs from 6 animals) and phtx larvae (red, n = 792 AZs from 14 NMJs from 5 animals) shown as box plots where horizontal lines represent median, boxes quartiles and whiskers 10^th^ and 90^th^ percentiles. Asterisks indicate significance level (* p < 0.05, ** p < 0.01, *** p < 0.001). (C) Scatter plots of type Ib AZs from a control animal (ctrl, left) and a GluRIIA receptor null mutant (GluRIIAko, right) representing chronic PHP. Localizations outside alpha shape are deleted for clarity. (D) Number of Brp localizations per AZ (p = 0.391), AZ area (p < 0.001) and Brp localization density (p < 0.001) for ctrl (grey, n = 872 AZs from 19 NMJs from 6 animals) and GluRIIAko larvae (blue, n = 1020 AZs from 19 NMJs from 6 animals). Scale bars 100 nm in (A) and (C).

Previous work discovered a proximo-distal functional and structural gradient with distal type Ib boutons containing more AZs, more Brp per AZ and releasing more glutamate than proximal boutons (Peled and Isacoff, 2011; Ehmann et al., 2014; Paul et al., 2015). To test whether the above described changes in phtx occur in a distinct spatial pattern within the NMJ we performed a subgroup analysis of AZs in boutons number 1-6 (Paul et al., 2015). Spearman correlation coefficients displayed no correlation between bouton number and Brp localization density in both groups (Supplementary Figure 2A). Thus, structural plasticity in acute homeostasis appears to occur homogeneously within the NMJ. Furthermore, there was neither a change in the total number of localizations nor the amount of unclustered Brp localizations outside AZs per bouton (Supplementary Figure 2B).

To clarify whether AZ compaction only appears after an acute homeostatic challenge or whether it is a general phenomenon in PHP, we used a knockout of the postsynaptic glutamate receptor subunit GluRIIA as a genetic, ‘chronic’ PHP model (GluRIIAko, Petersen et al., 1997; DiAntonio et al., 1999; Frank et al., 2006). We performed *d*STORM imaging and HDBSCAN analysis in GluRIIAko and wildtype larvae as described before (Figure 2C). Remarkably, we found comparable effects to PhTx treatment with unchanged Brp localization numbers per AZ, decreased AZ area and higher Brp density (Figure 2D). Thus, Brp scaffolds are also compacted in chronic PHP.

### AZ subcluster compaction

Next, we tested whether AZ reorganization during PHP also takes place within AZ subclusters (Figure 3A, C). Brp subcluster number per AZ was unchanged in phtx and GluRIIAko versus controls (Figure 3B, D). While Brp localization numbers in subclusters were unchanged, subcluster area was decreased (Figure 3B, D). Thus, mean Brp localization density in subclusters increased in acute and chronic PHP (Figure 3B, D). We also examined the radial distances between the c.o.m. of an individual AZ and the c.o.m.s of its subclusters and found that they decreased in phtx and GluRIIAko compared to controls (Figure 3B, D). In addition, the minimum distance between subcluster c.o.m.s was decreased in both forms of PHP, whereas the space between subclusters was only reduced in chronic PHP (Supplementary Figure 5). Taken together, these data show that Brp is compacted within AZ subclusters in acute and chronic homeostasis but the number of Brp molecules does not increase.

**Figure 3.**
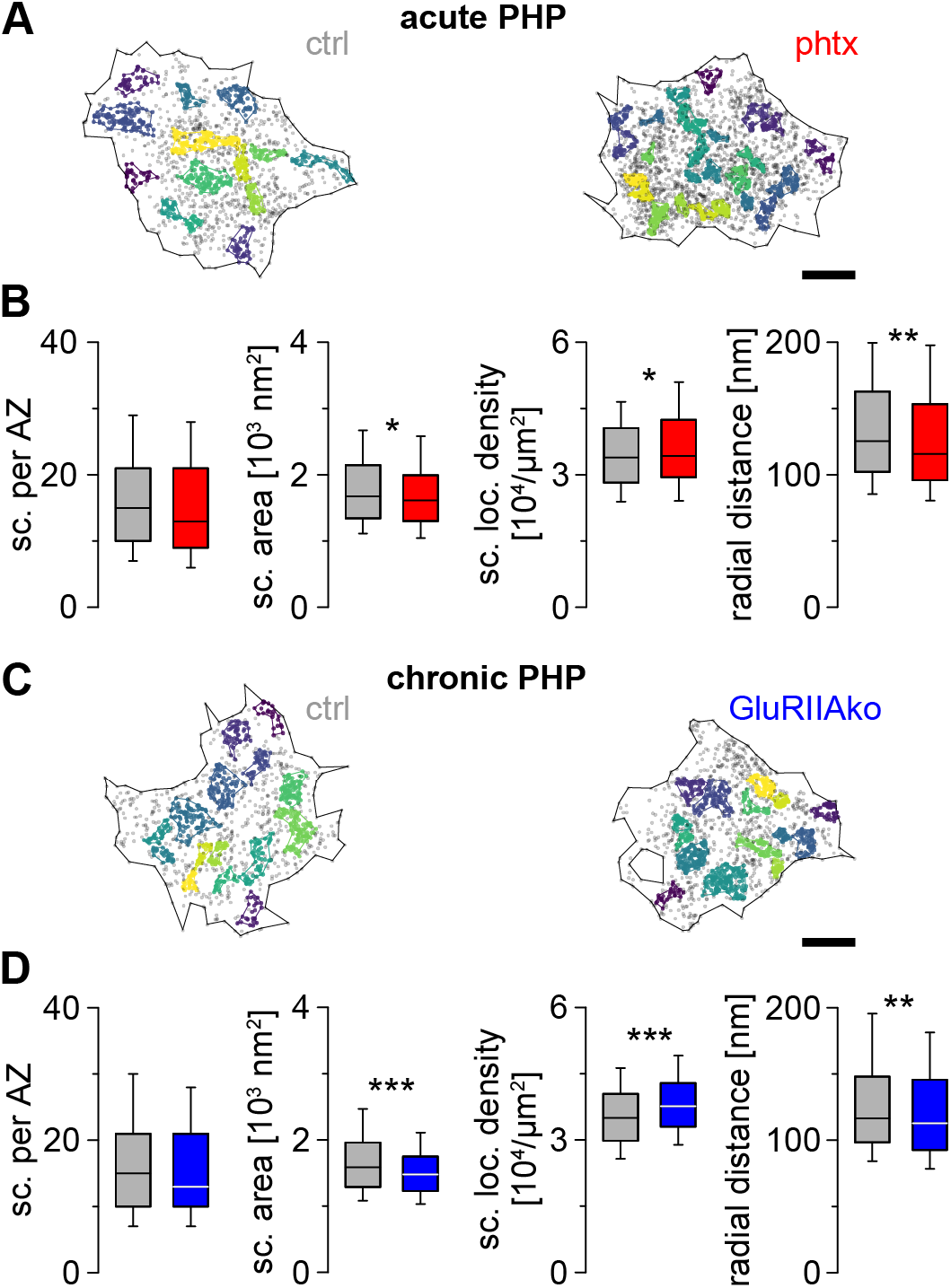
Homeostatic potentiation compacts AZ subclusters. (A) Scatter plots of type Ib AZs from a ctrl and a phtx animal. Brp subclusters are colored individually, unclustered localizations are displayed as grey dots. Localizations outside alpha shape are deleted for clarity. (B) Number of subclusters per AZ (p = 0.084), sc. area (p = 0.025), sc. localization density (p = 0.022) and radial distance between the AZ c.o.m. and the c.o.m. of individual subclusters (p = 0.003) for ctrl (grey, n = 568 AZs from 13 NMJs from 6 animals) and phtx larvae (red, n = 792 AZs from 14 NMJs from 5 animals). (C) Scatter plots of type Ib AZs from ctrl and GluRIIAko as shown in (A). (D) Number of subclusters per AZ (p = 0.118), sc. area (p < 0.001), sc. localization density (p < 0.001) and radial distance between the AZ c.o.m. and the c.o.m. of individual subclusters (p = 0.002) for ctrl (grey, n = 872 AZs from 19 NMJs from 6 animals) and GluRIIAko larvae (blue, n = 1020 AZs from 19 NMJs from 6 animals). Scale bars 100 nm in (A) and (C).

Next, we quantified the circularity of Brp clusters as the ratio of their Eigenvalues of the covariance matrix (ratio between 0 and 1 with 1 indicating a perfect circle, i.e. presumably in top view, see Material and Methods). AZ circularity was similar in phtx and control, thus, analysis of the entire dataset without orientation selection appears appropriate. We then investigated the relation between AZ area and circularity. Interestingly, this uncovered an inverse correlation (Supplementary Figure 2C). We interpret that low circularity is indicative of AZs in side view or of large Brp spots that partially arise from merged AZs lying nearby in 2D projection. The latter have been referred to as double ring structures, grouped CAZ units, or cluster AZs (Kittel et al., 2006; Ehmann et al., 2014; Akbergenova et al., 2018). Assuming that some structural parameters depend strongly on AZ orientation, we analyzed AZ area and radial distance in a subsample of AZs with circularity ≥ 0.5 and more planar orientation (Supplementary Figure 2D). Again, we found a pronounced decrease in AZ area and a similar decrease of radial distance in both groups. This indicates that AZ compaction in PHP appears regardless of AZ orientation.

In earlier studies abdominal muscle 4 which exclusively forms type Ib boutons was used to analyze structural effects of PhTx on AZs (Goel et al., 2017; Böhme et al., 2019). Therefore, we imaged PhTx treated AZs on abdominal muscle 4 and applied the described cluster analysis. This yielded similar results as shown for AZs on abdominal muscles 6/7 including subcluster parameters (Supplementary Figure 3).

### Correlative confocal-*d*STORM imaging of AZs

Earlier confocal and STED microscopy reported an increased fluorescence intensity in PHP and interpreted this as evidence for protein recruitment to the AZ (Weyhersmüller et al., 2011; Goel et al., 2017; Böhme et al., 2019). Since our results are in sharp contrast to these findings we wondered whether the difference in molecular density before and after PHP could bias AZ area quantification either in our localization-based approach or in intensity-based approaches (confocal and STED microscopy). To test this, we compared both methods using simulated circular clusters without noise and subclustering (Supplementary Figure 4). Our results indicate that the area of large objects with low molecular density (i.e. lower fluorescence intensity) can be grossly misjudged by pixel- and intensity-based quantification, which could explain the discrepancy between our and earlier findings. To further examine this experimentally, we perfomed sequential confocal-*d*STORM imaging of Brp in type Ib boutons (Figure 4A, B). Confocal data were analyzed with an intensity threshold of 100 a. u. (Figure 4Aii). Figure 4C shows enlarged AZs overlaid in confocal and *d*STORM resolution from Figure 4A and B in green, magenta and blue to mark AZ identity. The examples demonstrate area overestimation due to high localization density (Figure 4Ci) and show that on the other hand thresholding of confocal images can lead to complete loss of signal that is included in *d*STORM quantification (Figure 4Cii). Additionally, segmentation of closely spaced AZs is clearly superior using HDBSCAN-based quantification of localization data compared to thresholding-based quantification of confocal data (Figure 4Ciii). In a next step, we quantified AZs in the complete dataset after correlative confocal-*d*STORM imaging. Distribution of AZ area was broader in confocal microscopy than in *d*STORM (Figure 4D). As expected, we obtained positive correlations between the localization density per AZ measured with *d*STORM and the mean pixel intensity per AZ measured with confocal (Figure 4E). Interestingly, localization density correlated stronger with confocally measured than *d*STORM-measured AZ area (Figure 4F) matching our simulation results (Supplementary Figure 4). Confocal microscopy mostly overestimates AZ area, whereas underestimation only occurs at low AZ localization density (Figure 4G). This indicates that higher localization density in *d*STORM leads to an apparent enlargement of AZ area reported by thresholded confocal microscopy.

**Figure 4.**
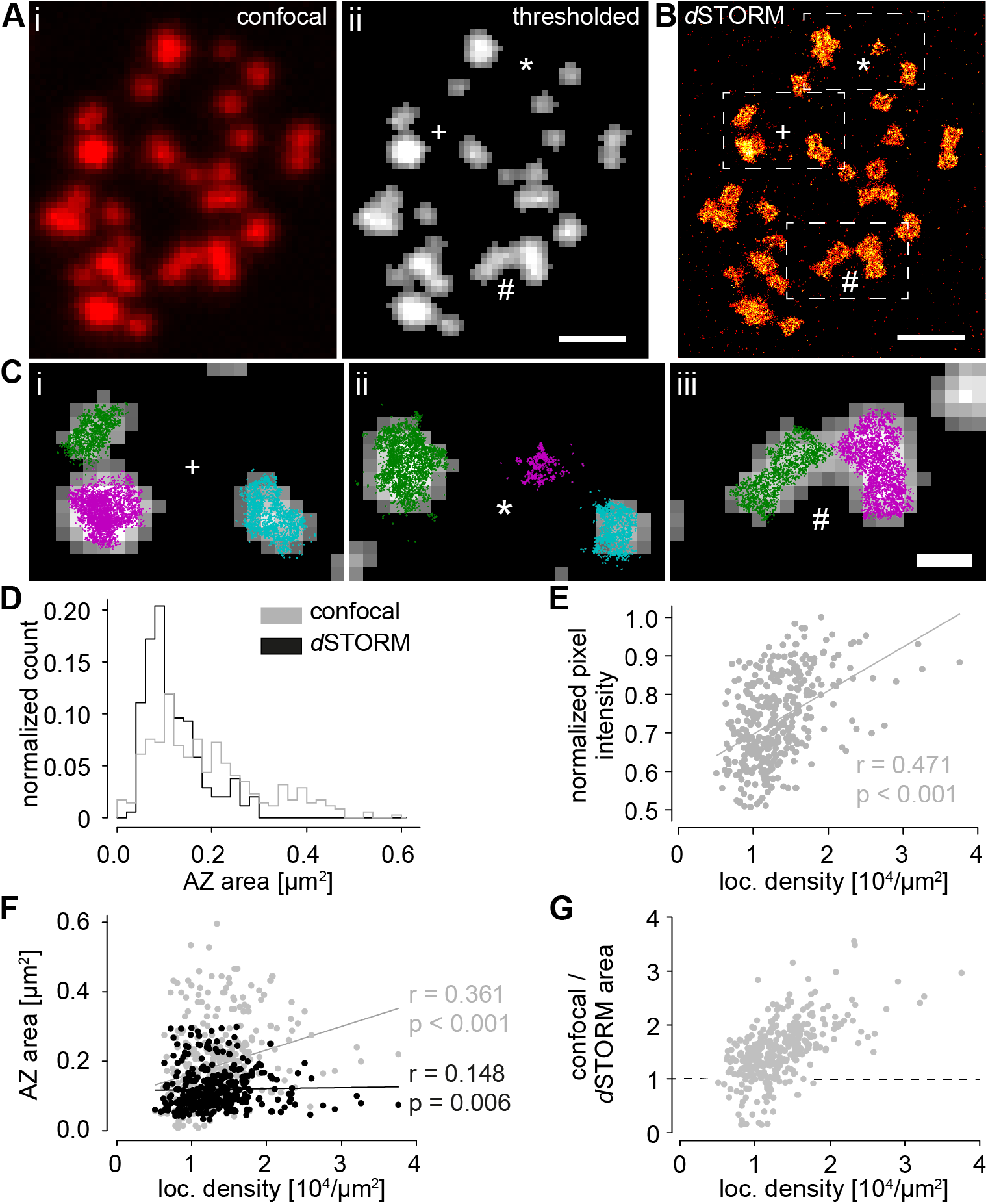
Correlative confocal-*d*STORM microscopy links increased localization density to increased confocal intensity and AZ area. (A) Confocal image of a type Ib bouton. (i) unprocessed original image and (ii) after applying a 100 a. u. threshold to the original image. Symbols mark corresponding regions in (B) enlarged in (C). (B) *d*STORM image of the same bouton as shown in (A). Boxes correspond to enlarged regions in (C). (C) Enlarged confocal AZs thresholded at 100 a. u. in grey corresponding to boxed regions in (B) and overlaid scatter plots of Brp localizations from *d*STORM (green, magenta, blue for cluster identity). Localization clusters at the edges are not displayed. Note that the green and magenta clusters in (Ci) are not separated in confocal microscopy after thresholding, that the magenta cluster in (Cii) is lost after confocal thresholding, and that the two clusters in (Ciii) are not separated by confocal microscopy. (D) AZ area in confocal (grey) and *d*STORM imaging (black, for D-F n = 343 corresponding AZs from 12 NMJs from 7 animals). (E) Spearman correlation coefficient shows a positive correlation between the normalized mean pixel intensity in confocal imaging and *d*STORM localization density. (F) Spearman correlation coefficients show a stronger correlation between AZ area in confocal imaging and localization density (grey) compared to AZ area measured with *d*STORM and localization density (black). (G) Ratio of confocal and *d*STORM area of corresponding AZs plotted against their localization density. Values below 1 (dashed black line) indicate an underestimation of confocal area quantification with respect to *d*STORM, whereas values larger than 1 indicate overestimation. Scale bars 1 μm in (A) and (B), 330 nm in (C).

### Homeostatic compaction is translated into an apparent area increase

In contrast to earlier studies (Weyhersmüller et al., 2011; Goel et al., 2017; Böhme et al., 2019) we found smaller AZ areas but increased localization density following PHP. To gain a better understanding of this discrepancy we performed simulations of confocal spatial resolution applying a Gaussian filter of 150 nm SD (Figure 5A, B). 3D histograms of two example AZs in control and phtx illustrate the applied thresholding (Figure 5C, compare Figure 4B in Weyhersmüller et al., 2011). In phtx, AZ area is decreased and localization density increased in original *d*STORM resolution (x-axes, Figure 5D, E), whereas both AZ area and mean pixel intensity increased in the same images in simulated confocal resolution (y-axes). We conclude that intensity-based quantification with confocal resolution may lead to inverse results concerning AZ area compared to localization-based analysis. In a last step, we simulated STED resolution as a less dramatic resolution decrease applying a Gaussian filter of 25 nm SD (Figure 5F). AZ area was measured similarly as mentioned above, AZ diameter was estimated from AZ area and intensity maxima per AZ were detected in selected planar-oriented AZs with a peak finding algorithm in FIJI after thresholding (Material and Methods; Böhme et al., 2019). This indicated an increase in AZ area, AZ diameter and number of intensity maxima per AZ after induction of PHP (Figure 5G), apparent findings contrary to our localization-based data evaluation. In summary, simulation of lower resolution gives inverse results for the number of clusters per AZ matching aforementioned findings.

**Figure 5.**
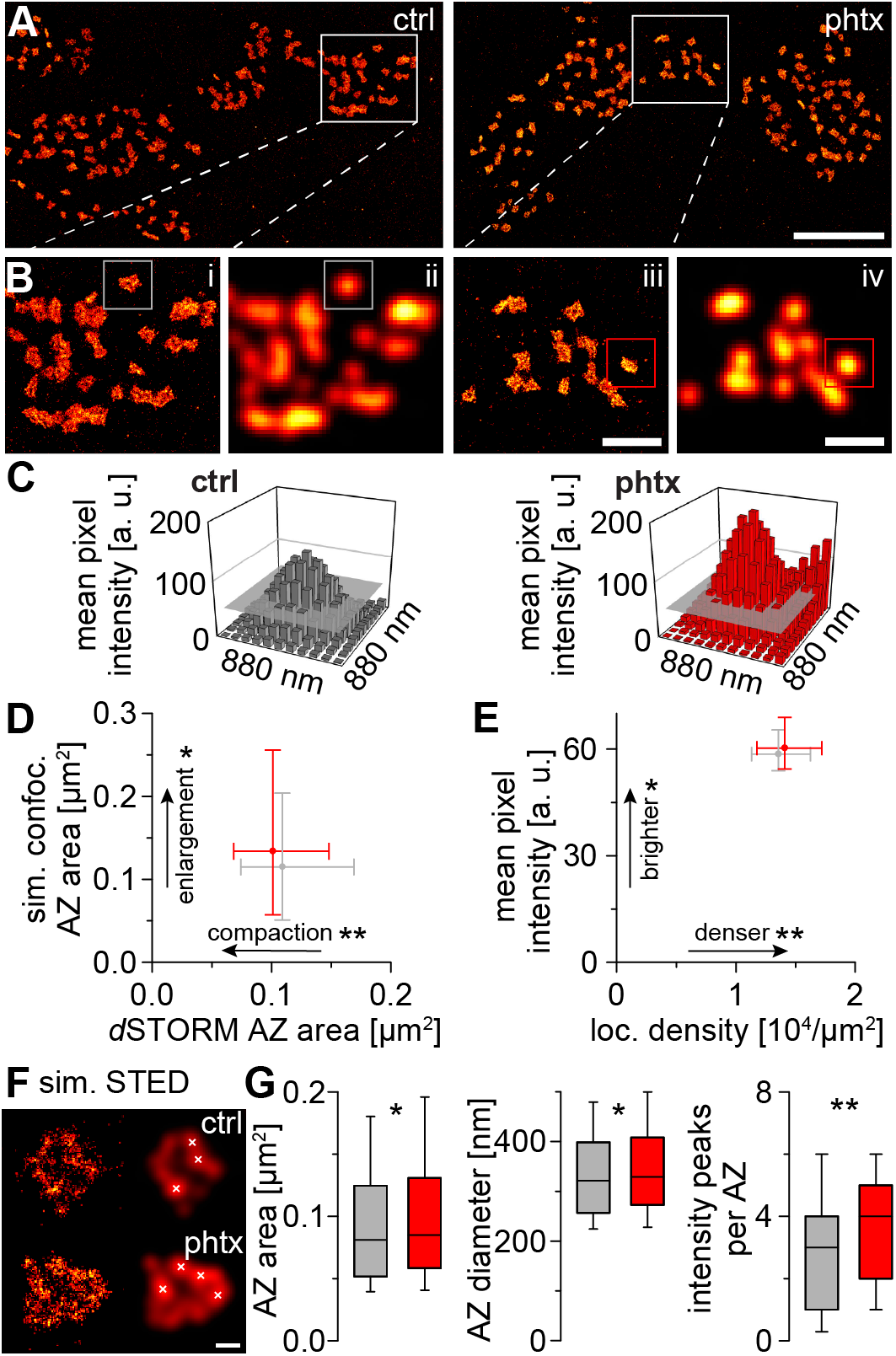
Compaction is reported as increased area in confocal simulation and as higher AZ diameter and more clusters in STED simulation. (A) Sections of *d*STORM images in ctrl and phtx (5 nm binning). Boxes highlight enlarged regions in (B). (B) Enlarged regions from (A) with *d*STORM resolution (i / iii) and simulated confocal resolution (ii / iv, 80 nm pixels, 150 nm Gaussian blur). Boxes highlight selected AZs in (C). (C) 3D bar plots of mean pixel intensity in confocal resolution of boxed AZs in (B). Grey plane indicates thresholding level of 50 a. u. which was used for quantification in (D, E). (D) AZ area (median (25^th^-75^th^ percentile)) quantified as shown in (C), i.e. by assuming confocal resolution, plotted against AZ area from original *d*STORM analysis (data from Figure 2B) in ctrl (grey, n = 345 AZs from 13 NMJs from 6 animals in confocal resolution for (D, E)) and phtx (red, n = 457 AZs from 14 NMJs from 5 animals in confocal resolution for (D, E), p = 0.026). Note the inverse, significant results for AZ area change. (E) Mean pixel intensity in simulated ‘confocal’ resolution (right, p = 0.016) plotted against *d*STORM localization density (data from Figure 2B) in ctrl and phtx. Also note the inverse, significant results for this analysis. (F) Example type Ib AZs of ctrl (upper panels) and phtx (lower panels) in *d*STORM resolution (left) and simulated STED resolution (25 nm Gaussian blur, right). White crosses indicate maxima detected with FIJI peak finding algorithm. (G) AZ area and AZ diameter, estimated under the assumption of a circular area, obtained from simulated STED analysis in ctrl (n = 591 AZs from 13 NMJs from 6 animals) and phtx (n = 820 AZs from 14 NMJs from 5 animals, p = 0.044 and 0.044) and number of intensity peaks, i.e. Brp modules, per AZ in ctrl (n = 122 AZs from 12 NMJs from 6 animals) and phtx (n = 186 AZs from 14 NMJs from 5 animals, p = 0.002). Scale bars 4 μm in (A), 1.2 μm in (B) and 100 nm in (F).

## DISCUSSION

We introduce AZ compaction as a mechanism for presynaptic structural adaptation in acute and chronic PHP. Compaction encompasses reduced AZ area and enhanced molecular density without increasing protein amount. Brp subclusters are likewise compacted and located closer to AZ centers during PHP. Notably, AZ compaction appears as an increase of AZ area in thresholding-based analysis, explaining the apparent contradiction with earlier results.

### Protein proximity promotes vesicle traffic

Despite substantial progress in single-molecule based microscopy, resolving the spatial relations between individual subcellular molecules remains challenging (Nahmani et al., 2017). In this study, we reason that *d*STORM and a localization-based clustering algorithm are a powerful experimental conjunction to decipher protein dense networks at single presynaptic AZs. The effects reported here are small in absolute terms, but very reproducible under different conditions (Figure 2 and 3, Supplementary Figure 2, 3 and 5). In the light of mechanistic considerations, it is plausible that changes of molecular proximities at AZs in the nanometer range lead to considerable functional consequences (Neher 1998). For example, it is established for the t-SNARE protein Syntaxin that the formation of protein clusters enhances local protein concentration, which in turn may raise the probability of protein interactions (Sieber et al., 2007; van den Bogaart et al., 2011). Furthermore, in hippocampal neurons RIM clusters with diameters of ~80 nm were described to show remarkable reorganization during synaptic plasticity (Tang et al., 2016). Molecular concentrations can be estimated from our data and Brp molecule numbers per AZ reported previously by Ehmann et al., 2014 assuming a distribution of the Brp^Nc82^ epitope in a hemisphere. For radii of 185 and 175 nm (Figure 2D, Supplementary Table 1) the hemisphere volumes before and after PHP are ~0.013 and 0.011 μm^3^. 170 Brp molecules (Ehmann et al., 2014) yield concentrations of 21 and 25 μM before and after PHP, respectively. Thus, the ~7 % density increase translates into a 19 % concentration increase. Brp subclusters are assumed to contain bundles of ~7 Brp molecules creating a platform for a single synaptic vesicle as described below (Ehmann et al., 2014). For hemispheres with subcluster radii of 22.5 and 21.7 nm (Figure 3D) this yields protein concentrations of 490 and 540 μM before and after PHP. While the above-mentioned estimates for Brp concentrations in the whole AZ are probably reasonably accurate, the estimates for the concentration in subclusters provide only a lower limit. Here, subcluster radii are most certainly overestimated due to the linkage error introduced by the combination of a primary antibody with the secondary F(ab’)_2_ fragment leading to a ‘labeling complex diameter’ of 26 nm (Ehmann et al., 2014). Thus, local protein concentrations could be even higher and, likewise, the difference in the two scenarios. This could be clarified in future studies using smaller labeling approaches, e.g. directly fluorophore-conjugated Brp^Nc82^ F(ab’)2 fragments. Brp was shown to directly or indirectly tether synaptic vesicles at AZs via its C-term (Hallermann et al., 2010; Scholz et al., 2019). Other coiled-coil proteins like Golgins tether and traffic vesicles at the Golgi network by guidance via multiple vesicle-binding sites (Gillingham and Munro, 2016; Cheung and Pfeffer, 2016). Likewise, Brp could provide a meshwork guiding vesicles via multiple vesicle-interaction sites represented by subclusters. The Brp^Nc82^ epitope is located close to the C-term and an increased protein concentration, achieved by AZ compaction, could enhance the probability of these interactions and facilitate vesicle supply. Taking the ~40.5 nm vesicle diameter from Karunanithi et al., 2002 and adding ~10 nm of protein trimmings (Takamori et al., 2006) resulting in 50-60 nm veritable vesicle extent, the distance between Brp clusters should fit approximately these dimensions. This is the case and since we observe compaction of Brp subclusters and a reduced radial distance this should further enhance trafficking of vesicles over the surface generated by Brp C-terms (Supplementary Figure 5). This structural adaption is in line with evidence for increased vesicle traffic during PHP (Weyhersmüller et al., 2011; Delvendahl et al., 2019).

Vesicle cluster formation depends on liquid-liquid phase separation, achieved by intrinsically disordered regions (IDRs, Milovanovic et al., 2018). Glutamine-rich IDRs were implicated in the formation of coiled-coils and protein agglomerates (Fiumara et al., 2010). The vesicle tethering Brp C-term comprises a stretch of glutamine residues (Hallermann et al., 2010). Vesicle traffic in liquid-liquid phase separation with Brp C-terms could profit from AZ compaction. While the molecular binding partners of the Brp C-term are still unclear, a functional interaction with Complexin was reported (Scholz et al., 2019). We assume that all Brp molecules forming one subcluster create a platform for tethering a single synaptic vesicle. This hypothesis matches reasonably well with EM studies describing on average 12 vesicles tethered to a single CAZ (Böhme et al., 2016) and with reconstructions of EM tomography data (Zhan et al., 2016).

### Compaction of the AZ scaffold without recruitment of Brp molecules

Localization data can be used to estimate protein numbers (Ehmann et al., 2014; Löschberger et al., 2012). In this study, the number of Brp localizations (i.e. Brp molecules) did not change after induction of homeostasis, which is in line with previous work ruling out protein synthesis during PHP (Frank et al., 2006). Confocal and STED microscopy provided data which were interpreted as evidence for protein incorporation into AZs during acute and chronic homeostasis (Weyhersmüller et al., 2011; Goel et al., 2017; Böhme et al., 2019). However, our data support an alternative interpretation (Figure 2B, D, Supplementary Figure 2B). Confocal and STED data were analyzed using pixel- and thresholding-based approaches that depend on signal intensity (Weyhersmüller et al., 2011; Goel et al., 2017; Böhme et al., 2019). Here, we show that large structures with low intensity appear smaller than small structures with higher intensity (Supplementary Figure 4C). We performed correlative confocal-*d*STORM imaging and demonstrated that a higher protein density in *d*STORM translates into an apparent area increase in confocal microscopy (Figure 4F). In confocal microscopy usually a stack of multiple slices is obtained, whereas in 2D-*d*STORM we sampled only signal from a single focal plane. However, we found no evidence for signal that is included in confocal images but is missing in corresponding *d*STORM data (compare Figure 4Ai and B). We measured that the z-range of boutons in confocal stacks ranges from 800 nm to 2 μm, roughly corresponding to the axial capture range of our *d*STORM measurements (Tokunaga et al., 2008). The correlation between localization density and AZ area in confocal microscopy is corroborated by simulating confocal resolution (Figure 5D, E).

### A molecular concept of homeostatic plasticity

Diverse cellular plasticity mechanisms enable the remarkable capacity of the nervous system to adapt, refine and increase in strength. Among them, the unique form of homeostatic plasticity stabilizes brain function on time scales of minutes to decades proceeding via different molecular pathways (Turrigiano et al., 2008; Younger et al., 2013; Orr et al., 2017; Ortega et al., 2018; Frank et al., 2020). We here report that compaction of the AZ scaffold is a crucial hallmark of homeostasis, due to the increase of protein density and therefore promotion of vesicle traffic. In contrast, disuse hypersensitivity, another form of synaptic plasticity induced by blockade of the AMPA glutamate receptor, was linked to AZ enlargement reversable by application of PhTx (Murthy et al., 2001; Thiagarajan et al., 2005). This illustrates that distinct forms of plasticity can exhibit varying structural expressions. At the *Drosophila* NMJ a proximo-distal gradient in release strength is accompanied by (ultra-)structural adaptations of AZs (Peled and Isacoff, 2011; Ehmann et al., 2014; Paul et al., 2015). To test whether this form of intraneuronal differentiation influences the ability of an AZ to undergo homeostatic plasticity, we analyzed our *d*STORM data according to this aspect. Interestingly, we find that PHP increases Brp density regardless of the bouton position (Supplementary Figure 2A).

We describe compaction on the level of individual AZs (i.e. clusters of Brp epitopes) and of Brp subclusters. However, the molecular mechanism of compaction for release strengthening might be extended to other AZ molecules. Orr et al., 2017 demonstrated that the Semaphorin2b-PlexinB complex regulates PHP via control of presynaptic actin. Whereas the role of actin for synaptic efficacy has been described unambiguously (Cingolani and Goda, 2008), the significance for compaction of AZs remains to be determined. Furthermore, AZ molecules located in membrane-proximity like ENaCs (Younger et al., 2013), presynaptic calcium channels (Turrigiano 2008), RIM and RIM-binding protein should be examined using superresolution microscopy to determine whether compaction is a universal plasticity concept during PHP. Finally, it appears promising to correlate transmission properties of individual AZs with its nanoarchitecture using transgenically expressed GCaMP Ca^2+^ sensors (Akbergenova et al., 2018; Gratz et al., 2019) to link synaptic activity with ultrastructure.

## MATERIAL AND METHODS

### Fly stocks

Flies were raised on standard cornmeal and molasses medium at 25 °C. *Drosophila melanogaster* male 3^rd^ instar larvae of the following strains were used for experiments: Wildtype: *w^1118^* (Bloomington *Drosophila* Stock Center). GluRIIA knockout: *DGluRIIA^AD9^/Df(2L)cl^h4^* (kindly provided by Mathias A. Böhme).

### Philanthotoxin treatment and and larval preparation

Philanthotoxin 433 tris (trifluoroacetate) salt (PhTx, P207 Sigma) was dissolved in dimethyl sulfoxide (DMSO) to obtain a stock solution of 4 mM and stored at −20 °C. For each experiment, the respective volume was further diluted with freshly prepared haemolymph-like solution (HL-3, Stewart et al., 1994) to a final PhTx concentration of 20 μM in 0.5 % DMSO. Control experiments were performed with the same DMSO concentration in HL-3. PhTx treatment of semi-intact preparations was performed essentially as described previously (Frank et al., 2006). In brief, larvae were pinned down in calcium-free, ice-cold HL-3 at the anterior and posterior endings, followed by a dorsal incision along the longitudinal axis. Larvae were incubated in 10 μl of 20 μM PhTx in DMSO for 10 minutes at room temperature (22 °C). Following this incubation time, PhTx was replaced by HL-3 and larval preparations were completed, followed by fixation and staining.

### Fixation, staining and immunofluorescence

After PhTx treatment and dissection, larvae were fixed with 4 % paraformaldehyde in phosphate buffered saline (PBS) for 10 minutes and blocked for 30 minutes with PBT (PBS containing 0.05 % Triton X-100, Sigma) including 5 % natural goat serum (Dianova). Primary antibodies were added for overnight staining at 4 °C. After two short and three long (20 min each) washing steps with PBT, preparations were incubated with secondary antibodies for 3 hours at room temperature, followed by two short and three long washing steps with PBT. Preparations were kept in PBS at 4 °C until imaging. All data were obtained from NMJs formed on abdominal muscles 6/7 in segments A2 and A3, except data for Supplementary Figure 3, which were obtained from NMJs formed on abdominal muscle 4 in segments A2-A4. Directly compared data (e.g. Figure 2B) were obtained from larvae stained in the same vial and measured in one imaging session.

### *d*STORM (*direct* stochastic optical reconstruction microscopy)

Super-resolution imaging of the specimen was performed essentially as previously reported (Ehmann et al., 2014; Paul et al., 2015). Preparations were incubated with monoclonal antibody (mAb) Brp^Nc82^ (1:100, Antibody Registry ID: AB_2314866, Developmental Studies Hybridoma Bank) and secondary antibody goat α-mouse F(ab’)_2_ fragments labelled with Alexa Fluor647 (1:500, A21237, Thermofisher). Boutons were visualized using Alexa Fluor488 conjugated goat α-horseradish-peroxidase antibody (α-hrp, 1:250, Jackson Immuno Research). After staining, larval preparations were incubated in 100 mM mercaptoethylamine (MEA, Sigma-Aldrich) in a 0.2 M sodium phosphate buffer, pH 7.8 to 7.9, to allow reversible switching of single fluorophores during data acquisiton (van de Linde et al., 2008). The buffer additionally included an oxygen-scavenging system (10% (wt/vol) glucose, 10 U/ml glucose oxidase and 200 U/ml catalase). In all experiments, images were acquired using an inverted microscope (Olympus IX-71, 60x, NA 1.49, oil immersion) equipped with a nosepiece-stage (IX2-NPS, Olympus). 647 nm (F-04306-113, MBP Communications Inc.) and 488 nm (iBEAM-SMART-488-S, Toptica) lasers were used for excitation of Alexa Fluor647 and Alexa Fluor488, respectively. Laser beams were passed through clean-up filters (BrightLine HC 642/10, Semrock and ZET 488/10, Chroma, respectively), combined by a dichroic mirror (LaserMUX BS 473-491R, 1064R, F38-M03, AHF Analysentechnik), and directed onto the probe by an excitation dichroic mirror (HC Quadband BS R405/488/532/635, F73-832, AHF Analysentechnik). The emitted fluorescence was filtered with a quadband-filter (HC-quadband 446/523/600/677, Semrock) and a longpass- (Edge Basic 635, Semrock) or bandpass-filter (Brightline HC 582/75, Semrock) for the red and green channel, respectively, and divided onto two cameras (iXon Ultra DU-897-U, Andor) using a dichroic mirror (HC-BS 640 imaging, Semrock). The green channel was used for visualizing individual presynaptic boutons in normal fluorescence microscopy. For the red channel, image resolution was 126 nm x 126 nm per pixel to obtain super-resolution of Brp. Single fluorophores were localized and high resolution-images were reconstructed with rapi*d*STORM (Heilemann et al., 2008; van de Linde et al., 2011; Wolter et al., 2010; Wolter et al., 2012; www.super-resolution.de). Only fluorescence spots with more than 12000 photons were analyzed and subpixel binning of 10 nm px^−1^ was applied. For visualization of representative *d*STORM measurements (Figure 4B, Figure 5A, B, F), reconstructed images from rapi*d*STORM with 5 nm binning were opened in FIJI

(Schindelin et al., 2012) and where contrast enhanced for clarity. Localization precision was determined with the NeNa algorithm (nearest neighbor based analysis, Endesfelder et al. 2014), implemented in the LAMA software package (LocAlization Microscopy Analyzer; Malkusch and Heilemann, 2016) in 108 *d*STORM measurements of Brp and was 6.3 ± 0.6 nm (mean ± SD) in this study. To check for homogenous illumination of the samples we analyzed and compared localization intensity of all *d*STORM measurements, (i) for the whole image, and (ii) only for analyzed regions and found no differences in A/D counts between the groups.

### Confocal microscopy

For confocal imaging larvae were mounted in PBS and imaged using a commercial confocal laser scanning microscope (Zeiss LSM 700) equipped with an oil-immersion objective (Plan-Apochromat 63x/1.40 Oil M27). Alexa Fluor647 and Alexa Fluor488 were excited with the 639 nm and the 488 nm diode laser lines, repectively. The microscope was controlled via ZEN 12 software (black edition, Zeiss AG). For each z-stack 7-13 slices with 200 nm axial spacing were obtained. The gain of the photomultiplier tubes was adjusted to 700 and 600 V and laser power was set to 1.5 % and 1 % in the red and green channel, respectively, to obtain a good signal with little photobleaching allowing subsequent *d*STORM imaging. The pinhole was set to 70.6 μm, corresponding to 1.32 Airy units in the red and 1.73 Airy units in the green channel. Images were recorded in 1024 x 1024 (lines x pixels) format with 16-bit data depth. Given an approximate resolution of 200 nm in the red channel, the pixel size, adjusting the zoom factor to 1.2, was set to about 83 nm for sufficient data sampling. Settings resulted in a pixel dwell time of 3.15 μs.

### Data evaluation

Localization microscopy data were analyzed with custom written Python code (https://www.python.org/, language version 3.6) and the web-based Python interface Jupyter (https://jupyter.org/index.html). Localization tables from rapi*d*STORM were directly loaded and analyzed. Prior to the Python-based analysis the regions of interest (ROI) were masked in the reconstructed, binned images from rapi*d*STORM using FIJI (1.440, Schindelin et al., 2012). These ROIs corresponded to the terminal 6 boutons according to the α-hrp staining. In earlier work on super-resolved Brp-data, density-based spatial clustering of applications with noise (DBSCAN) was used to identify AZ subclusters (Ehmann et al., 2014). Given the parameters k and ε, DBSCAN considers a group of localizations as a cluster if there are at least k residues in a circle with radius ε around a certain localization (Ester et al., 1996). Here, we used an improved approach called hierarchical density-based spatial clustering of applications with noise (HDBSCAN) that, in contrast, extracts the most robust clusters from a cluster hierarchy over varying ε environments that are least sensible to ε variation, i.e. have the longest lifetime in the cluster tree (Campello et al., 2013). The algorithm is thus more powerful for cluster detection in data with variable density. We used the Python implementation of HDBSCAN (McInnes et al., 2017; https://github.com/scikit-learn-contrib/hdbscan), which takes ‘minimum cluster size’ and ‘minimum samples’ as the main free parameters, and performed two steps of clustering with different parameters on our localization data: (i) to identify the AZs, and (ii) to extract the subclusters from the AZs. Setting minimum samples to smaller values than minimum cluster size allows the algorithm to be less conservative, i.e. extract clusters that might be smaller than minimum cluster size but very robust in the cluster hierarchy. We explored optimal clustering parameters for the detection of active zones and varied minimum cluster size and minimum samples in a certain range (Figure 1B, Supplementary Figure 1). Too high values falsely merge adjacent AZs together (Figure 1C) whereas too low values lead to fragmenting AZs into smaller clusters (Figure 1E). A wide range of parameters delivered robust results and all Brp clusters in this study were extracted with the combination 100 and 25 for minimum cluster size and minimum samples. A second HDBSCAN on the individual AZ clusters was performed with minimum cluster size 24 and minimum samples 6, and a similar number of subclusters with a comparable spatial extent compared to the DBSCAN-based subcluster analysis of Ehmann et al., 2014 was found (compare Figure 1H for subcluster radii and Figure 3B, D for subcluster areas). For subcluster analysis, the cluster selection method was changed to ‘leaf’ clustering, which comprises a tendency to more homogenous clusters by extracting those that lie on leaf nodes of the cluster tree rather than the most stable clusters. To quantify cluster area, we computed 2D alpha shapes using CGAL (Computational Geometry Algorithms Library, https://www.cgal.org) in Python. The geometrical concept of alpha shapes can be used to calculate the shape and thus the area of a set of points. Given a finite set of points and an alpha value α, an edge of the shape is drawn between two points whenever they lie on the boundary of a disk with radius α that does not contain any other point (Edelsbrunner and Mücke, 1994). To get the alpha shapes of the AZ clusters and AZ subclusters we choose α-values of 800 nm^2^ and 300 nm^2^, respectively. The subcluster center of mass (c.o.m.) was calculated as mean of the x- and y-values of all localizations of the subcluster, and the AZ c.o.m. as mean of its subcluster c.o.m.s. To estimate the distance of subcluster c.o.m. from AZ c.o.m. the Euclidean distance of these points was computed. The distance to the nearest subcluster c.o.m. was determined for each individual subcluster c.o.m. and the mean value per AZ was obtained (Supplementary Figure 5A). To get the distance between subclusters the mean subcluster radius per AZ was subtracted from the subcluster center distance (Supplementary Figure 5B). For evaluation of Brp cluster circularity, the ratio of the Eigenvalues of each cluster was computed, where 1 indicates a perfect circle and values < 1 indicate decreasing circularity (Supplementary Figure 2C). Exclusion criteria for outliers in all *d*STORM data evaluations of Brp were AZ area < 0.03 μm^2^ (Ehmann et al., 2014) and > 0.3 μm^2^, absolute localization counts per AZ > 8000 and mean AZ localization density > 60000 localizations per μm^2^ (about 3-5-fold median). Additional exclusion criteria for type Ib neuron recordings of Brp were average AZ localization count < 1000 and at the same time average AZ area < 0.095 μm^2^ per image, indicative of insufficient data quality. For visualization of the localization-based results (Figure 1A, C-E, G, Figure 2A, C, Figure 3A, C, Figure 4C, Supplementary Figure 3A) scatter plots were created in Python. The H function (Figure 1F) as derivative of Ripley’s K function was computed using Python package Astropy (Robitaille et al., 2013) for each individual AZ and for the random Poisson distribution. Curves for display were averaged (mean ± SD). The function was evaluated in nm steps for radii from 0 to 120 nm and without correction for edge effects. Simulated localization data (Supplementary Figure 4) for a given radius r were created in Python using the function numpy.random.uniform() passing the arguments -r and r. Thus, random numbers >= -r and < r were created and used as x- and y-coordinates of random points. Simulation was further constrained by computing the Euclidean distance of the simulated point and the center of the simulated point cloud (x = 0, y = 0) and discarding all points with a distance > r. This gave rise to point clouds consisting of 1500 points with circular distributions in a circle with radius r around the center. For HDBSCAN analysis nine random point clouds per given radius in the same coordinate system were used as input spacing the center points of adjacent point clouds three radii apart. To simulate confocal resolution the point coordinates were binned into 5 nm pixels. Binned images were imported as text images in FIJI and automatically converted to 32-bit images were raw integrated density is equal to the absolute number of localizations in a pixel. A Gaussian filter with 150 nm standard deviation (SD) was applied to blur the images. The maximally possible pixel intensity after blurring was 0.265 arbitrary units (a. u.) corresponding to 1500 points in one pixel before blurring. To achieve a somewhat realistic gain all images were equally scaled to a maximum of 0.35 a. u. Thresholding was performed in 8-bit with 50 a. u.

Confocal images were processed and evaluated in FIJI. Z-stacks were maximum projected and Brp spots segmented with a pixel intensity threshold of 100 a. u. in 8-bit using the ‘Analyze Particles’ function (Figure 4Aii). Mean pixel intensity of resulting masks was measured in the original images. Corresponding Brp spots, extracted with 100 a. u. threshold, and Brp localization clusters were assigned manually. Only clusters that were clearly distinguishable in both confocal and *d*STORM without confluence to neighboring signal were used. Bouton height ranged from 0.8 to 2 μm in confocal imaging. To ensure comparability to earlier quantification approaches we created maximum projections of confocal data. The capture range of our *d*STORM measurements is at least 1 μm, but probably even higher (Tokunaga et al., 2008). To minimize bias arising from analysis of different focal planes we inspected all correlative images and found no evidence for signal that was included in confocal stacks but missing in *d*STORM images (compare Figure 4Ai and B). Additionally, only AZs were correlated that could be clearly assigned in both imaging techniques.

To simulate confocal resolution (Figure 5A, B) localization tables from rapi*d*STORM with 5 nm binning were converted to FIJI-readable density matrices where raw integrated density corresponds to the localization count in a pixel. Images were contrast enhanced to obtain 0.1 % saturated pixels, a Gaussian filter with 150 nm SD was applied and the images were scaled to a pixel size of 80 nm. The same images as for the localization-based analysis were included and only pixels in the same ROIs as described above were used for subsequent processing and analysis. Images were converted to 8-bit and the gain was adjusted by scaling all pixels in both images to the same value by setting the brightest of all analyzed pixels in all images to the maximum value of 255 a. u.. Following a standard protocol for the quantification of confocal data of the *Drosophila* NMJ (Schmid et al., 2008; Weyhersmüller et al., 2011) a minimal threshold of 50 a. u. was applied, and the ‘Analyze Particles’ function in FIJI was used to create individual masks from the thresholded images. The resulting masks were applied to the non-thresholded simulated confocal images to quantify AZ area as well as the mean intensity of AZs. STED resolution in Figure 5F was simulated similarly, but a Gaussian filter with 25 nm SD was applied to the density matrices. The gain of all images was adjusted by setting 175 a. u. to the maximum. AZ area and mean pixel intensity per AZ were measured with a threshold of 50 a. u. (Weyhersmüller et al., 2011) and only Brp spots with an area > 0.03 μm^2^ were analyzed. The diameter of Brp spots was calculated under the assumption of a circular area. To quantify intensity maxima per AZ planar-oriented, ring-like AZs were selected. Selection was performed blinded with respect to the experimental groups. Single AZs were cut out from the whole images. Thresholding was performed with 100 a. u. and all pixels below the threshold were removed. The maxima were extracted using the ‘Find Maxima’ function in FIJI (Prominence = 0) and quantified per AZ.

### Statistics

Statistical analyses were performed with Sigma Plot 13 (Systat Software). Shapiro-Wilk was used to test normality. If data were not normally distributed, we used the non-parametric Mann-Whitney rank sum test for statistical analysis and reported data as median (25^th^-75^th^ percentile). If data were normally distributed they were reported as mean ± SD unless indicated otherwise (Figure 1F). Asterisks indicate the significance level (* p < 0.05, ** p < 0.01, *** p < 0.001) and n denotes sample number. In box plots, horizontal lines represent median, boxes quartiles and whiskers 10^th^ and 90^th^ percentiles. Scatter plots show individual data points unless indicated otherwise. Bin counts in histograms were normalized to the total number of observed events which was set to 1. Color codes in contour plots (Figure 1B, Supplementary Figure 1) were logarithmically scaled. Linear regression curves were fitted and Spearman correlation coefficient r and statistical significance p of correlations were evaluated in Sigma Plot. All plots were produced with Sigma Plot software and figures assembled using Adobe Illustrator (Adobe, 2015.1.1 release). Supplementary Table 1 contains all numerical values not stated in text and figure legends including p-values and samples sizes.

### Code and data availability

The authors declare that custom written Python code and all data sets supporting the findings of this work are available from the corresponding authors.

## ACKNOWLEDGEMENTS

This work was supported by grants from the German Research Foundation (DFG) TRR166/B06 to A.L.S. and M.H., TRR166/B02 to S.D., TRR166/B04 to M.S., KI1460/5-1 and KI1460/4-1 to R.J.K. and from the IZKF Würzburg to A.L.S. and M.H. (N229). The authors thank T. Langenhan for scientific discussions and M. Oppmann and F. Köhler for technical assistance. The authors declare no conflict of interest.

## SUPPLEMENTARY MATERIAL

**Supplementary Figure 1.**
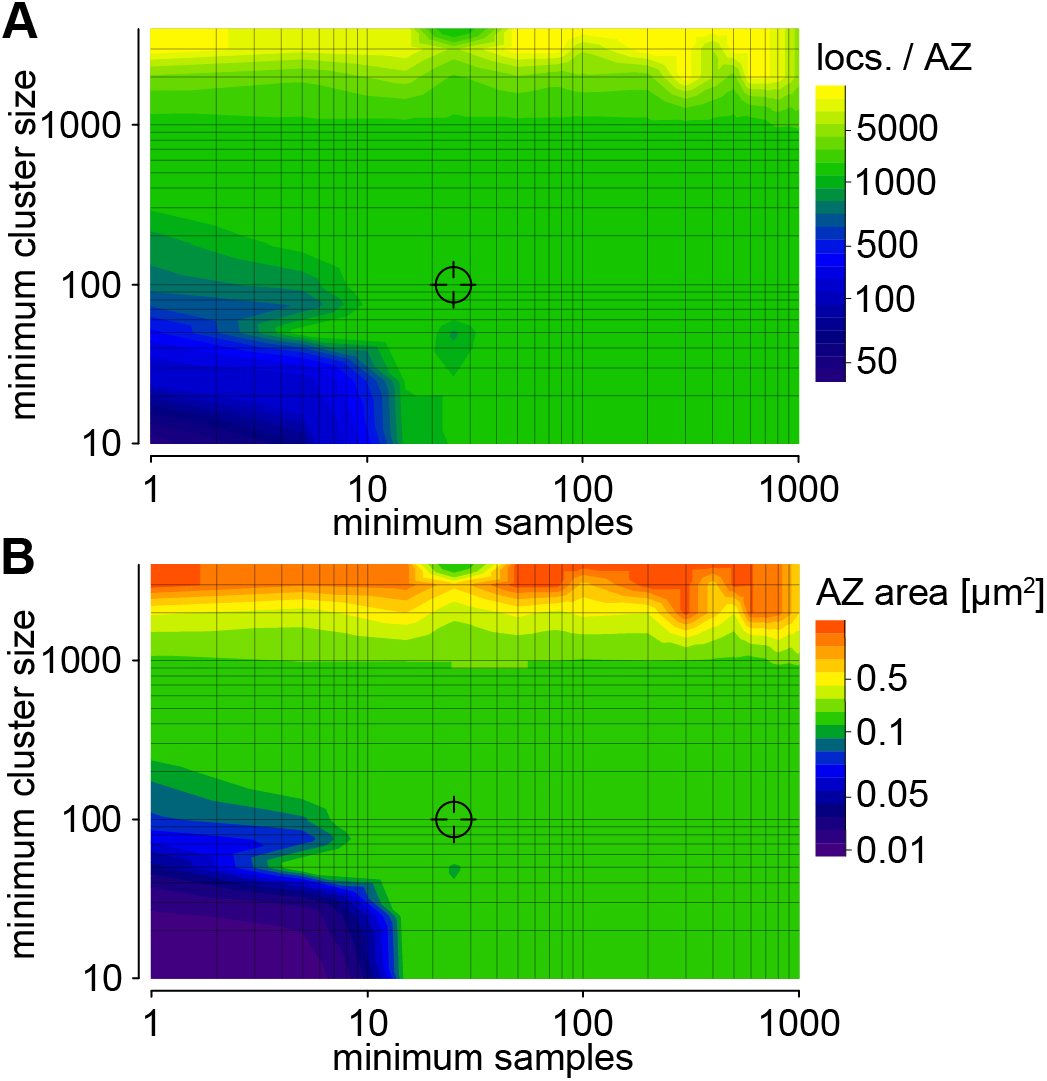
Exploring HDBSCAN parameters. *Related to Figure 1*. Contour plots of the mean number of localizations per AZ (A) and the mean AZ area (B) for the data shown in Figure 1A depending on the HDBSCAN clustering parameters ‘minimum samples’ and ‘minimum cluster size’. The parameter combination used for analysis throughout this study is indicated (minimum cluster size 100, minimum samples 25).

**Supplementary Figure 2.**
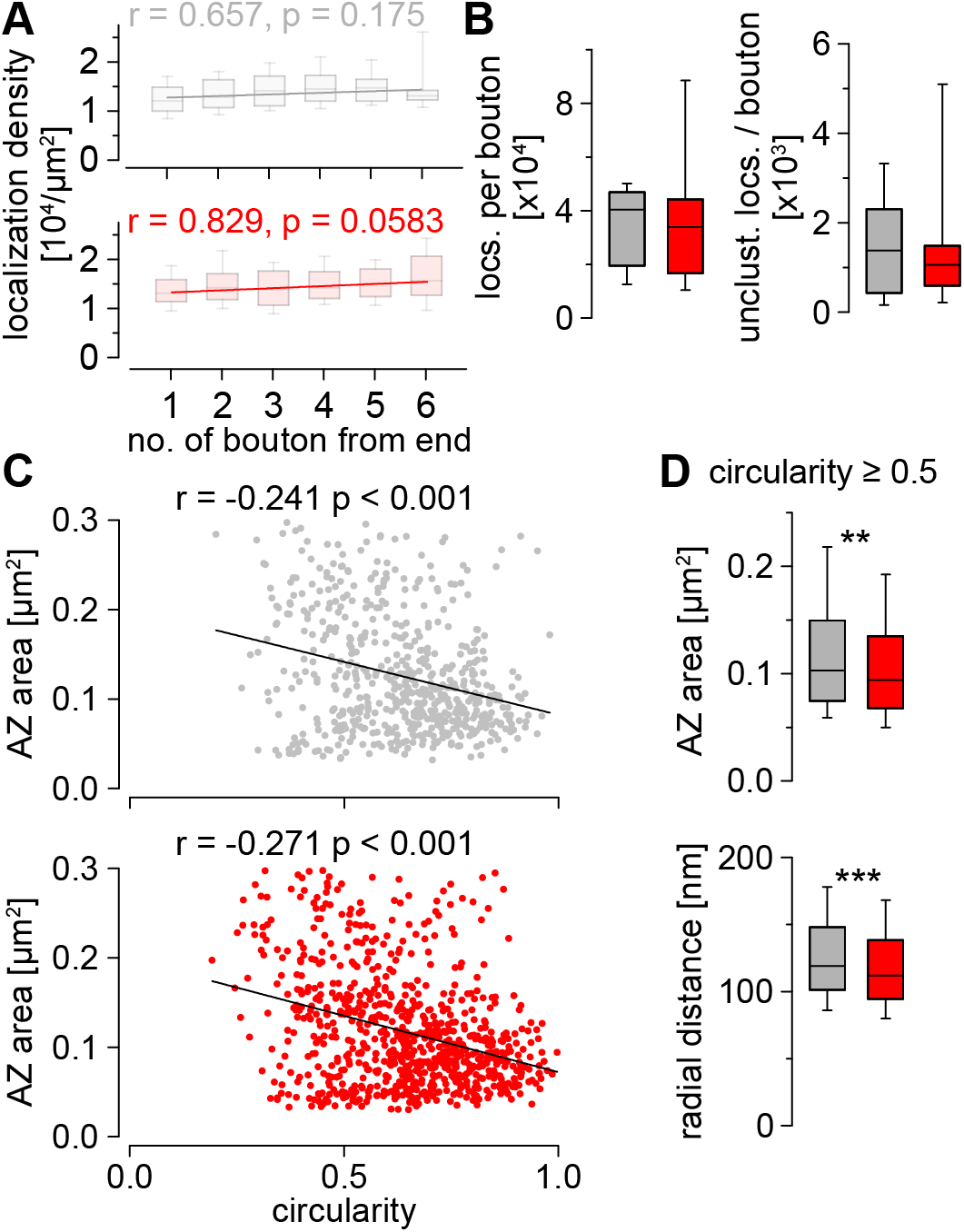
Spatial differentiation, bouton analysis and circularity. *Related to Figures 2 and 3*. (A) Brp localization density shown as boxplots for individual boutons number 1-6 from the end of the bouton string. Spearman correlation coefficients show no significant correlation between AZ localization density and bouton position in ctrl (grey, n = 240, 140, 102, 71, 54, 19 AZs for boutons 1-6) and phtx (red, n = 251, 134, 146, 117, 78, 51 AZs for boutons 1-6, respectively). (B) Number of all Brp localizations per bouton and number of unclustered (unclust.) localizations per bouton in ctrl (n = 13 distal Ib boutons from 6 animals) and phtx (n = 14 distal Ib boutons from 5 animals, p = 0.577 and 0.903). (C) Scatter plots illustrating correlation of AZ circularity and area in ctrl and phtx and linear regression lines. Spearman coefficients r and p-values show negative correlations in both groups. (D) AZs with circularity ≥ 0.5 (indicative for an AZ in top view) were used to compare AZ organization in ctrl and phtx. Box plots of AZ area and of the radial distance between the AZ c.o.m. and the c.o.m. of individual subclusters (n = 442 AZs from 13 NMJs from 6 animals and n = 621 AZs from 14 NMJs from 5 animals in ctrl and phtx, respectively, p = 0.002 and < 0.001).

**Supplementary Figure 3.**
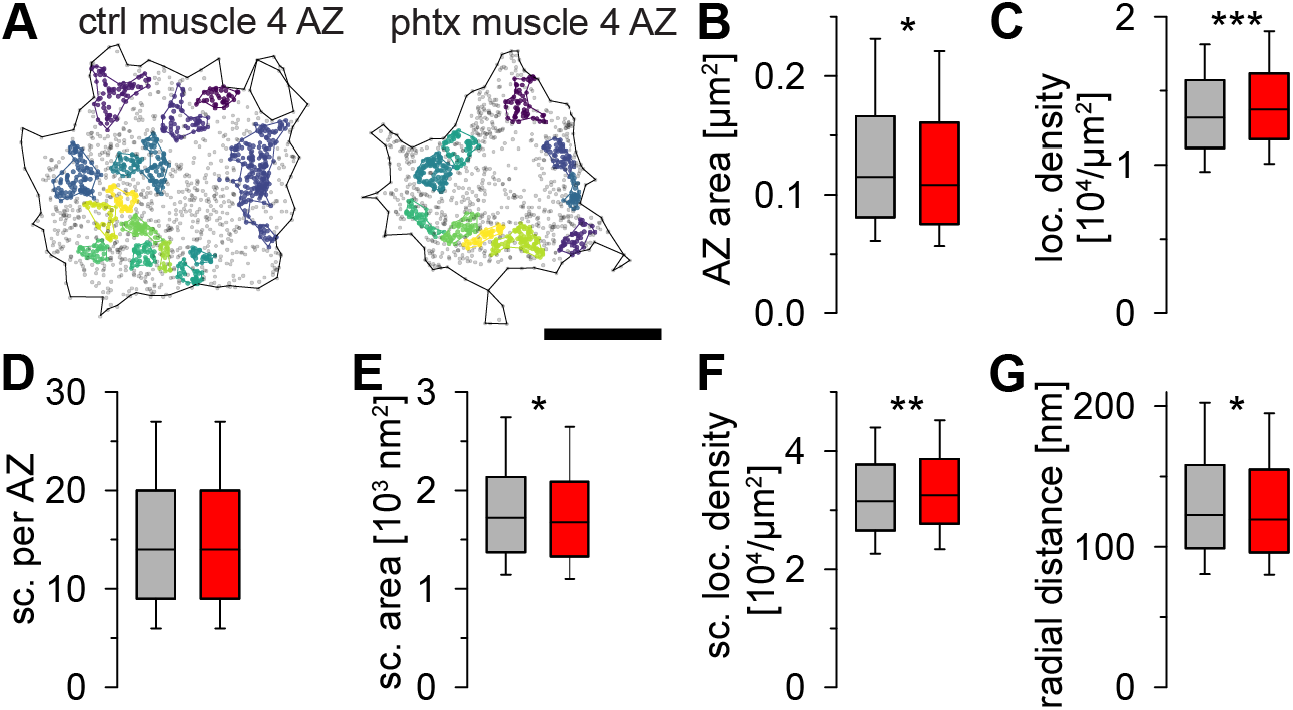
AZ nanoarchitecture in type Ib boutons from NMJs on abdominal muscle 4 of *Drosophila* 3^rd^ instar larvae. *Related to Figures 2 and 3*. (A) Scatter plots of a ctrl and a phtx type Ib AZ on muscle 4. (B, C) AZ area (B, p = 0.018) and Brp localization density (C, p < 0.001) in ctrl (grey, n = 1183 AZs from 20 NMJs from 6 animals for B-G) and phtx (red, n = 1445 AZs from 22 NMJs from 8 animals for B-G). (D-G) Subcluster numbers per individual type Ib AZ (D, p = 0.803), subcluster area (E, p = 0.020), subcluster localization density (F, p = 0.003) and radial distance between the AZ c.o.m. and the c.o.m. of individual subclusters (G, p = 0.022) in both groups. Scale bar 200 nm in (A).

**Supplementary Figure 4.**
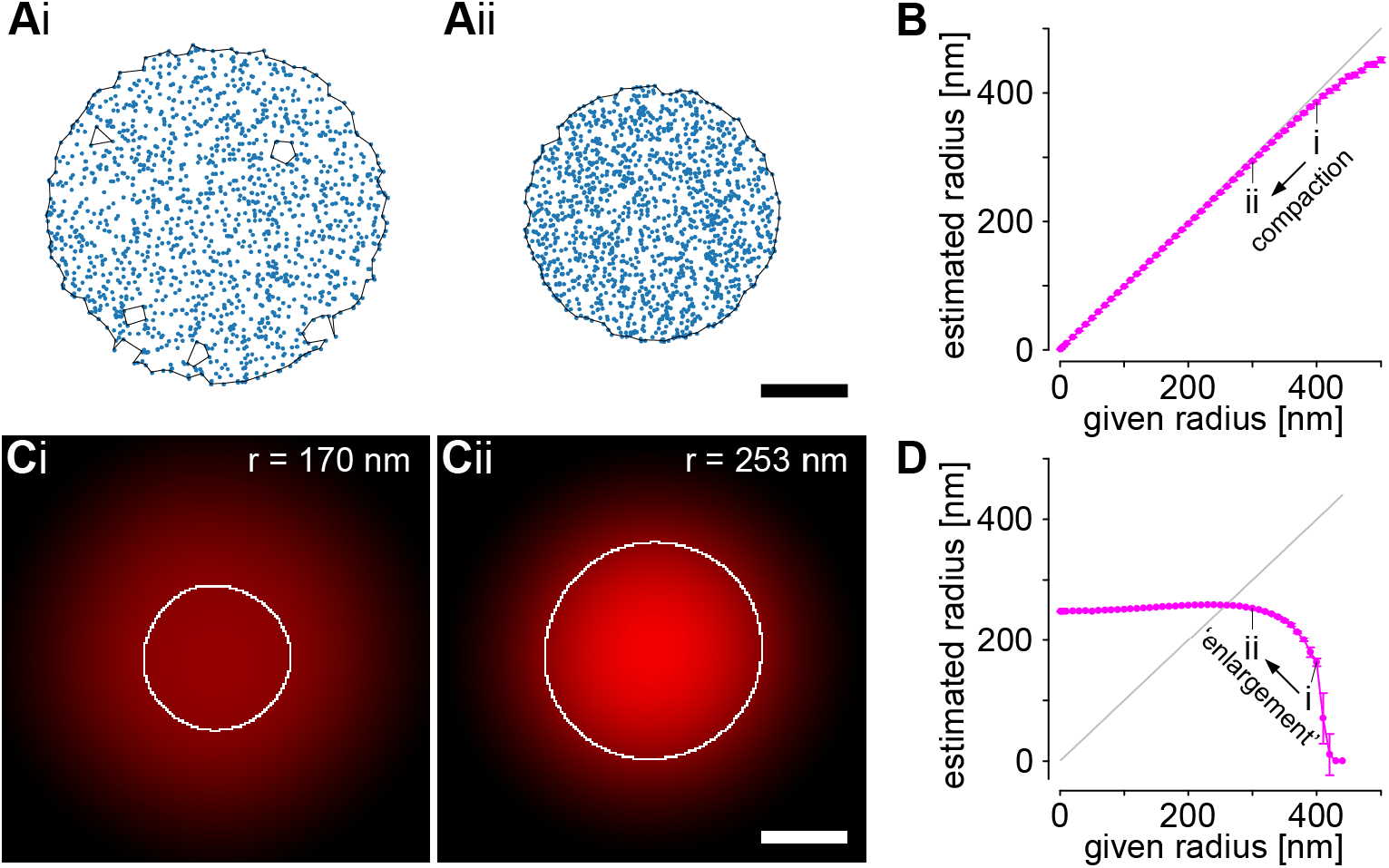
Limits of area quantification. (A) Simulation of 1500 random localizations (blue, compare ctrl value in Figure 2B, D) randomly distributed in circular areas with radius (i) 400 nm and (ii) 300 nm. Black lines indicate alpha shapes for area quantification. Holes in Ai exemplify the local failure of alpha shape-based area quantification. (B) Radius of simulated localization clusters (magenta, mean ± SD, n = 9 simulated clusters per given radius) estimated from alpha shape area under the assumption of a circular area and line for real radius (grey, equal to given radius) plotted against the given radius for simulation. Symbols (i, ii) correspond to the given radii used in (A). This revealed a robust area quantification and allowed us to define a critical density of about 2700 localizations per μm^2^ above which the estimated radius was always at least 95 % of the given radius. The first given radius below the critical density was 430 nm which exceeds the observed maximal AZ radius. We conclude that localization density is not limiting for area quantification given the high densities of AZs and of AZ subclusters in our data. (C) Data from (A) after binning in 5 nm pixels, application of a Gaussian filter of 150 nm SD and intensity scaling to impose confocal resolution. White lines indicate a 50 a. u. threshold used for area quantification (compare standard procedures for quantification of confocal data: Schmidt et al., 2008; Weyhersmüller et al., 2011; Goel et al., 2017; Böhme et al., 2019). Radii estimated from the area under the assumption of a circular area are displayed. Strikingly, the radius measured in confocal resolution by thresholding shows an inverse effect compared to the given radii and the radii computed by alpha shapes. (D) Radii after imposing confocal resolution and thresholding on simulated localization clusters (magenta, mean ± SD, n = 10 simulated clusters per given radius) and line for real radius (grey) plotted against the given radius for simulation. Symbols (i, ii) correspond to the given radii used in (C). Note the drop of the estimated radius above a given radius of 250 nm (corresponding to a localization density roughly below 7600 locs. per μm^2^). This implies that the area of large objects with low molecular density (i.e. lower fluorescence intensity) is systematically underestimated by pixel- and intensity-based quantification offering an explanation for the apparent discrepancy between our data and earlier findings. Scale bars 200 nm in (A) and (C).

**Supplementary Figure 5.**
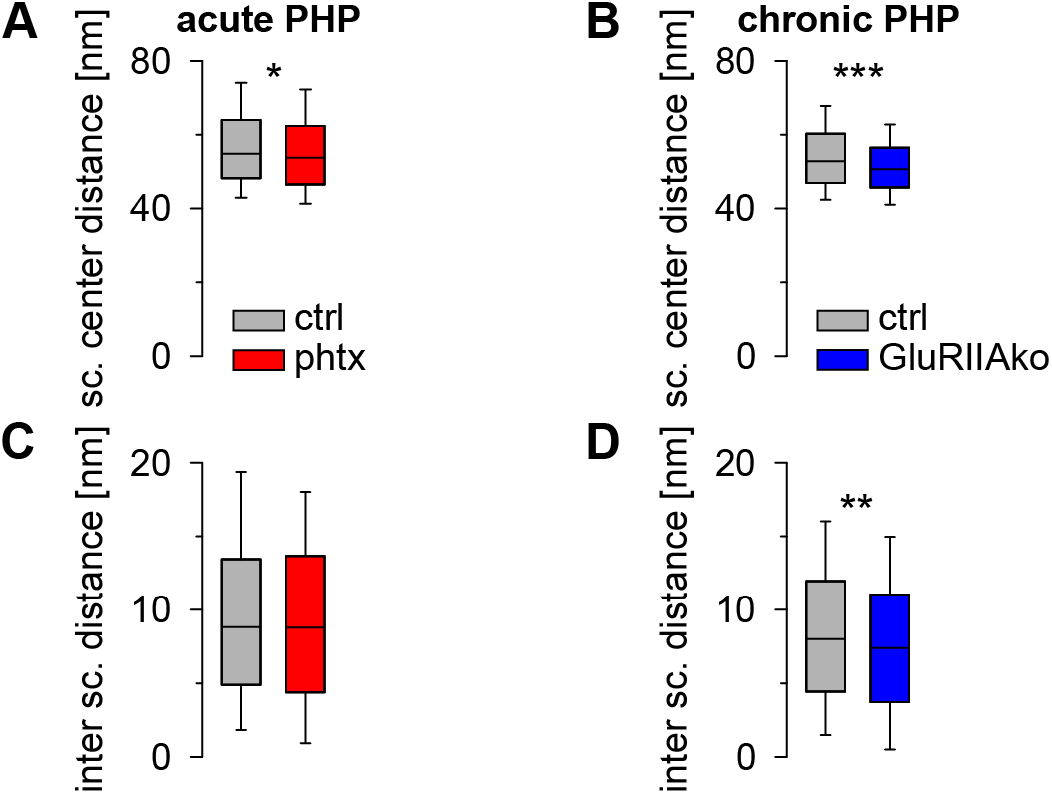
Homeostatic potentiation reduces subcluster spacing. (A, B) Minimum distance between subcluster centers (mean per AZ). Sc. center distance is decreased in acute PHP (ctrl: grey, n = 568 AZs from 13 NMJs from 6 animals for (A, C); phtx: red, n = 790 AZs from 14 NMJs and 5 animals for (A, C); p = 0.024) and in chronic PHP (ctrl: grey, n = 872 AZs from 19 NMJs from 6 animals for (B, D); GluRIIAko: blue, n = 1020 AZs from 19 NMJs from 6 animals for (B, D); p < 0.001). (C, D) Distance between subclusters (mean sc. diameter per AZ substracted from sc. center distance) in acute (C) and chronic (D) PHP. The distance between subclusters (inter sc. distance) is unchanged in acute PHP (p = 0.354) but reduced in chronic PHP (p = 0.007).

**Supplementary Table 1.**
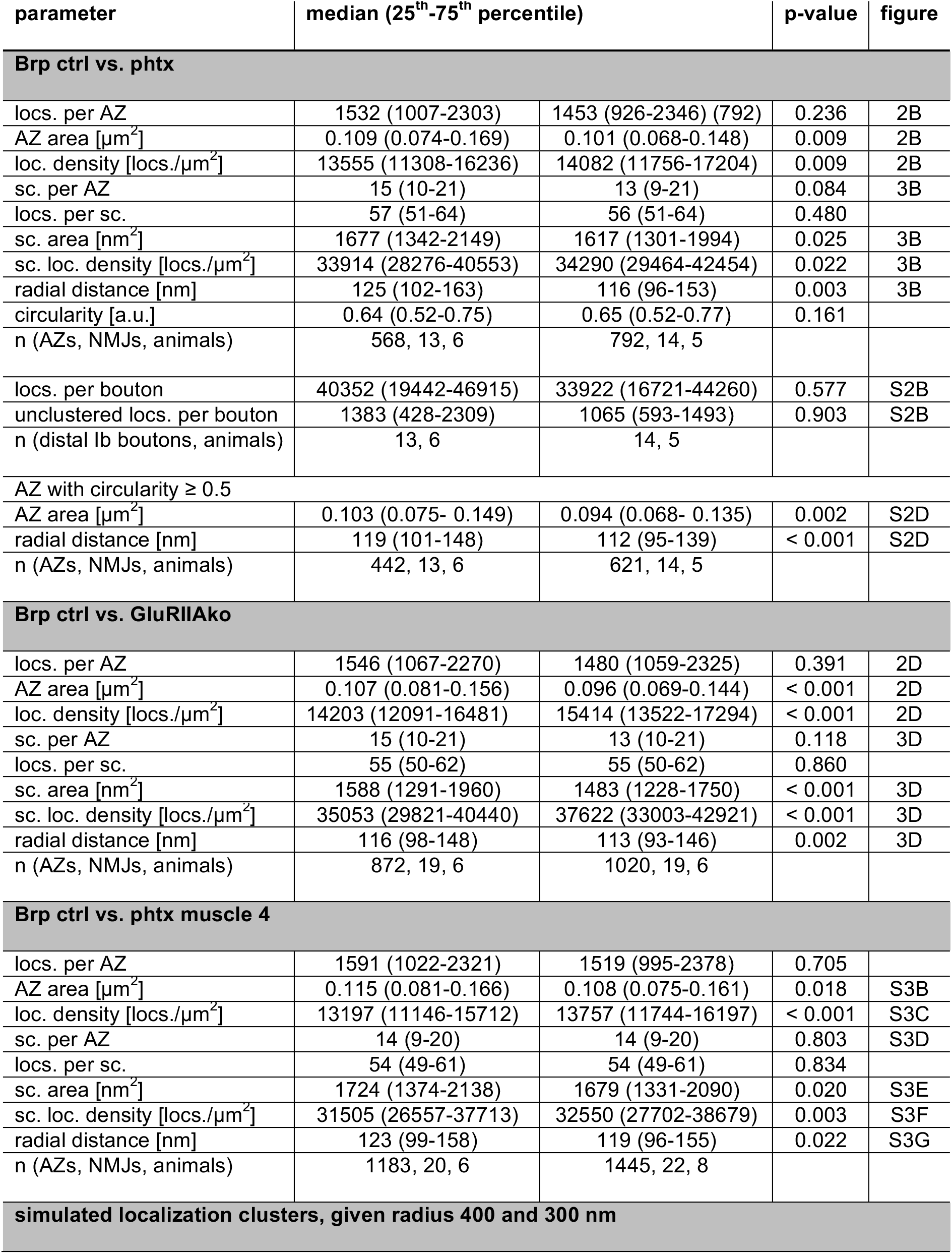
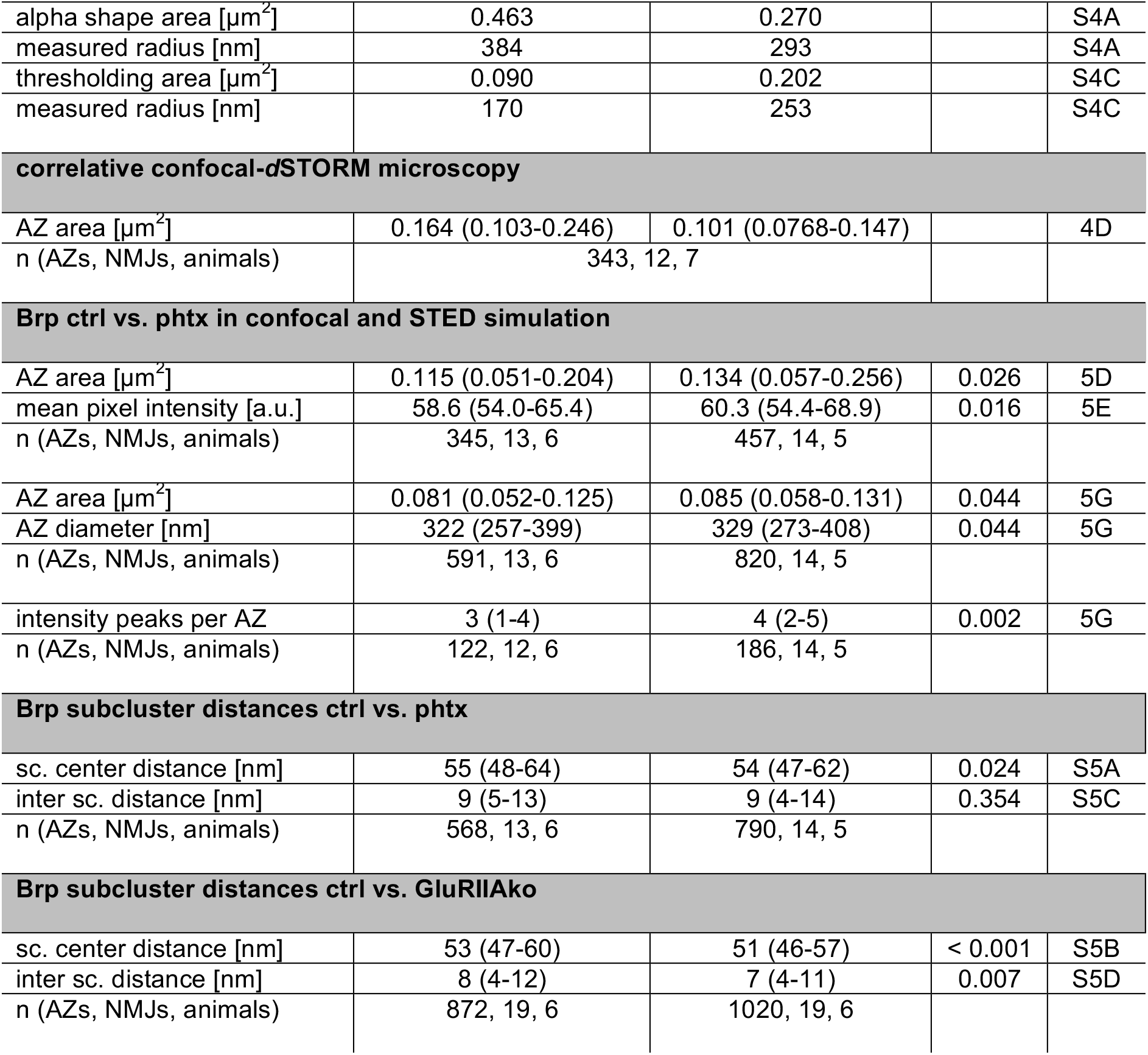
Data summary and statistical information. Numerical values not stated in text and figure legends including p-values and sample sizes. Data were not normally distributed and Mann-Whitney rank sum test was used to evaluate statistical significance. The largest number stated for n indicates the sample size used for the statistical test.

| parameter | median (25<sup>th</sup>-75<sup>th</sup> percentile) |  | p-value | figure |
| --- | --- | --- | --- | --- |
| Brp ctrl vs. phtx |  |  |  |  |
| locs. per AZ | 1532 (1007-2303) | 1453 (926-2346) (792) | 0.236 | 2B |
| AZ area [μm<sup>2</sup>] | 0.109 (0.074-0.169) | 0.101 (0.068-0.148) | 0.009 | 2B |
| loc. density [locs./μm<sup>2</sup>] | 13555 (11308-16236) | 14082 (11756-17204) | 0.009 | 2B |
| sc. per AZ | 15 (10-21) | 13 (9-21) | 0.084 | 3B |
| locs. per sc. | 57 (51-64) | 56 (51-64) | 0.480 |  |
| sc. area [nm<sup>2</sup>] | 1677 (1342-2149) | 1617 (1301-1994) | 0.025 | 3B |
| sc. loc. density [locs./μm<sup>2</sup>] | 33914 (28276-40553) | 34290 (29464-42454) | 0.022 | 3B |
| radial distance [nm] | 125 (102-163) | 116 (96-153) | 0.003 | 3B |
| circularity [a.u.] | 0.64 (0.52-0.75) | 0.65 (0.52-0.77) | 0.161 |  |
| n (AZs, NMJs, animals) | 568, 13, 6 | 792, 14, 5 |  |  |
| locs. per bouton | 40352 (19442-46915) | 33922 (16721-44260) | 0.577 | S2B |
| unclustered locs. per bouton | 1383 (428-2309) | 1065 (593-1493) | 0.903 | S2B |
| n (distal lb boutons, animals) | 13, 6 | 14, 5 |  |  |
| AZ with circularity ≥ 0.5 |  |  |  |  |
| AZ area [μm<sup>2</sup>] | 0.103 (0.075- 0.149) | 0.094 (0.068- 0.135) | 0.002 | S2D |
| radial distance [nm] | 119 (101-148) | 112 (95-139) | < 0.001 | S2D |
| n (AZs, NMJs, animals) | 442, 13, 6 | 621, 14, 5 |  |  |
| Brp ctrl vs. GluRIIAko |  |  |  |  |
| locs. per AZ | 1546 (1067-2270) | 1480 (1059-2325) | 0.391 | 2D |
| AZ area [μm<sup>2</sup>] | 0.107 (0.081-0.156) | 0.096 (0.069-0.144) | < 0.001 | 2D |
| loc. density [locs./μm<sup>2</sup>] | 14203 (12091-16481) | 15414 (13522-17294) | < 0.001 | 2D |
| sc. per AZ | 15 (10-21) | 13 (10-21) | 0.118 | 3D |
| locs. per sc. | 55 (50-62) | 55 (50-62) | 0.860 |  |
| sc. area [nm<sup>2</sup>] | 1588 (1291-1960) | 1483 (1228-1750) | < 0.001 | 3D |
| sc. loc. density [locs./μm<sup>2</sup>] | 35053 (29821-40440) | 37622 (33003-42921) | < 0.001 | 3D |
| radial distance [nm] | 116 (98-148) | 113 (93-146) | 0.002 | 3D |
| n (AZs, NMJs, animals) | 872, 19, 6 | 1020, 19, 6 |  |  |
| Brp ctrl vs. phtx muscle 4 |  |  |  |  |
| locs. per AZ | 1591 (1022-2321) | 1519 (995-2378) | 0.705 |  |
| AZ area [μm<sup>2</sup>] | 0.115 (0.081-0.166) | 0.108 (0.075-0.161) | 0.018 | S3B |
| loc. density [locs./μm<sup>2</sup>] | 13197 (11146-15712) | 13757 (11744-16197) | < 0.001 | S3C |
| sc. per AZ | 14 (9-20) | 14 (9-20) | 0.803 | S3D |
| locs. per sc. | 54 (49-61) | 54 (49-61) | 0.834 |  |
| sc. area [nm<sup>2</sup>] | 1724 (1374-2138) | 1679 (1331-2090) | 0.020 | S3E |
| sc. loc. density [locs./μm<sup>2</sup>] | 31505 (26557-37713) | 32550 (27702-38679) | 0.003 | S3F |
| radial distance [nm] | 123 (99-158) | 119 (96-155) | 0.022 | S3G |
| n (AZs, NMJs, animals) | 1183, 20, 6 | 1445, 22, 8 |  |  |
| simulated localization clusters, given radius 400 and 300 nm |  |  |  |  |
| alpha shape area [$\mu\text{m}^2$] | 0.463 | 0.270 | | S4A |
| measured radius [nm] | 384 | 293 |  | S4A |
| thresholding area [$\mu\text{m}^2$] | 0.090 | 0.202 | | S4C |
| measured radius [nm] | 170 | 253 |  | S4C |
| correlative confocal-dSTORM microscopy |  |  |  |  |
| AZ area [$\mu\text{m}^2$] | 0.164 (0.103-0.246) | 0.101 (0.0768-0.147) | | 4D |
| n (AZs, NMJs, animals) | 343, 12, 7 |  |  |  |
| Brp ctrl vs. phtx in confocal and STED simulation |  |  |  |  |
| AZ area [$\mu\text{m}^2$] | 0.115 (0.051-0.204) | 0.134 (0.057-0.256) | 0.026 | 5D |
| mean pixel intensity [a.u.] | 58.6 (54.0-65.4) | 60.3 (54.4-68.9) | 0.016 | 5E |
| n (AZs, NMJs, animals) | 345, 13, 6 | 457, 14, 5 |  |  |
| AZ area [$\mu\text{m}^2$] | 0.081 (0.052-0.125) | 0.085 (0.058-0.131) | 0.044 | 5G |
| AZ diameter [nm] | 322 (257-399) | 329 (273-408) | 0.044 | 5G |
| n (AZs, NMJs, animals) | 591, 13, 6 | 820, 14, 5 |  |  |
| intensity peaks per AZ | 3 (1-4) | 4 (2-5) | 0.002 | 5G |
| n (AZs, NMJs, animals) | 122, 12, 6 | 186, 14, 5 |  |  |
| Brp subcluster distances ctrl vs. phtx |  |  |  |  |
| sc. center distance [nm] | 55 (48-64) | 54 (47-62) | 0.024 | S5A |
| inter sc. distance [nm] | 9 (5-13) | 9 (4-14) | 0.354 | S5C |
| n (AZs, NMJs, animals) | 568, 13, 6 | 790, 14, 5 |  |  |
| Brp subcluster distances ctrl vs. GluRIIAko |  |  |  |  |
| sc. center distance [nm] | 53 (47-60) | 51 (46-57) | < 0.001 | S5B |
| inter sc. distance [nm] | 8 (4-12) | 7 (4-11) | 0.007 | S5D |
| n (AZs, NMJs, animals) | 872, 19, 6 | 1020, 19, 6 |  |  |

